# A pathogen effector FOLD diversified in symbiotic fungi

**DOI:** 10.1101/2022.12.16.520752

**Authors:** Albin Teulet, Clement Quan, Edouard Evangelisti, Alan Wanke, Weibing Yang, Sebastian Schornack

## Abstract

Pathogenic fungi use secreted effector proteins to suppress immunity and support their infection, but effectors have also been reported from fungi that engage in nutritional symbioses with plants. Sequence based effector comparisons between pathogens and symbiotic arbuscular mycorrhizal (AM) fungi are hampered by the huge diversity of effector sequences even within closely related microbes.

Here we used a systematic protein structure modelling approach to classify the secretome of the AM fungus *Rhizophagus irregularis*. We identified secreted proteins with high structural similarity to *Fusarium oxysporum* f. sp. *lycopersici* dual domain (FOLD) effectors, which occur in low numbers in fungal pathogen genomes. Contrastingly, genes encoding FOLD proteins from AM fungi (MycFOLDs) are found in enlarged and diversified gene families.

Our structure-model comparison suggests that MycFOLDs are similar to carbohydrate binding motifs. Different MycFOLD genes are expressed during colonisation of different hosts and *MycFOLD-17* transcripts accumulate in plant intracellular arbuscules.

The exclusive presence of MycFOLDs across unrelated plant-colonising fungi, their inducible expression, lineage specific sequence diversification, and transcripts in arbuscules support the hypothesis that FOLD proteins act as effectors during plant colonisation by symbiotic and pathogenic fungi.

## Introduction

Most land plants engage with fungi to form nutritional symbioses, but fungi also constitute important plant pathogens causing devastating diseases. Successful plant colonisation often relies on microbe-secreted effector proteins which operate in the host to suppress immunity, support fungal entry, and ensure fungal sustenance. To date, most effectors have been characterised from plant pathogenic fungi where they had initially been studied due to their activity to trigger the plant immune system in host plants carrying cognate intracellular disease resistance genes or surface receptors. Such effectors have been termed Avirulence (Avr) genes.

Importantly, effectors have also been predicted from genomes of symbiotic fungi (Miyauchi *et al*, 2020, Lahrmann *et al*. 2015, Perotto *et al*, 2018) including arbuscular mycorrhizal (AM) Glomeromycotina fungi such as *Rhizophagus irregularis. R. irregularis* SP7 (Kloppholz *et al*., 2011) and RiNLE1 (Wang *et al*., 2021) suppress plant immune responses. The lysin motif (LysM) effector RiSLM (Zeng *et al*., 2019) binds fungal chitin thus masking the fungus from the plant immune system. The strigolactone induced effector RiSIS (Tsuzuki *et al*., 2016) is required for full colonisation.

To date, all protein crystal structures of effectors from plant-associated microbes originate from pathogens. An emerging theme is the occurrence of effector families defined by a common structure. Examples are the RXLR-WY (e.g. AVR3a) or RXLR-LWY (e.g. PSR2) effector families of oomycetes (Boutemy *et al*., 2011, He *et al*., 2019), the MAX fold effectors from *Magnaporthe oryzae* (AVR-Pia, AVR1-CO39, AvrPiz-t; Zhang *et al*., 2013; Guo *et al*., 2018; Varden *et al*., 2019) and *Pyrenophora tritici-repentis* (e.g. ToxB; Nyarko *et al*., 2014), the LARS effectors of *Cladosporium fulvum* (now *Fulvia fulva*) and *Leptosphaeria maculans* (e.g. Ecp11-1, AvrLm4-7; Lazar *et al*., 2022), the RALPH effectors from *Blumeria graminis* (e.g. BEC1054; Pennington *et al*., 2019), ToxA family from *Pyrenophora tritici-repentis* and *Fusarium oxysporum* f. sp. *lycopersici* (e.g ToxA and Avr2; Sarma *et al*., 2005, Di *et al*., 2017) and finally the *F. oxysporum* f. sp. *lycopersici* (*Fo*l) dual domain (FOLD) effectors (e.g. FoAvr1, FoAvr3, FoSix6; Yu *et al*, 2022).

Members of the FOLD effector family belong to the secreted-in-xylem (Six) group of *Fusarium* proteins (Yu *et al*., 2022). The effectors Avr1 (Six4) and Avr3 (Six1) trigger an immune response in tomato carrying the cognate I and I-3 immune receptors and their N-terminal domain is sufficient for this avirulence activity. Six6 and Six13, two additional members of the FOLD family, were identified using structure model prediction and homologs are found in many pathotypes of *F. oxysporum* and other fungal genera including *Colletotrichum, Ustilaginoidea, Leptosphaeria, Pyrenophora* and *Bipolaris* (Yu *et al*., 2022). Six6 contributes to virulence and suppresses effector-triggered immunity (Gawehns *et al*., 2014).

Effectors are under constant evolutionary pressure to diversify their amino acid sequence to evade perception by the host immune system. This limits their genome-wide identification across divergent fungal species. However, effectors with low sequence similarity often retain a conserved protein structure. The rise of fast, *ab initio* protein structure prediction algorithms such as DeepMind’s AlphaFold2 has enabled proteome-wide predictions and the classification of proteins based on their structure rather than sequence (Bordin *et al*., 2022; Akdel *et al*., 2022). However, no structural model-based classification of the secreted proteins from endomycorrhizal AM fungi including their effectors has been reported. The extent to which AM fungi harbour proteins with structural similarity to known pathogen effector families therefore remains unknown.

AM fungi from the Glomeromycotina clade represent important symbiotic partners for plants to gain access to additional nutrients. Although AM fungal species differ in their morphology and colonisation structures (Dickson, 2004), they display similar colonisation strategies. AM fungi enter plant roots by means of a hyphopodium and subsequently grow inter- and sometimes intracellular hyphae which can differentiate into intracellular arbuscules in root cortex cells. The arbuscule is a fungal structure surrounded by a periarbuscular matrix and bordered by a carbohydrate-rich periarbuscular membrane towards the living host cell (Balestrini and Bonfante, 2014).

Here, we report a structural model-based classification of all secreted proteins of the AM symbiotic fungus *R. irregularis*. A comparison of the structural models with fungal pathogen effectors identified the FOLD group of effectors are shared between *Fol* and *R. irregularis*. While FOLD effectors occur in pathogenic fungal genomes in one to three copies, this family is expanded in some AM fungal genomes. The N-termini of secreted *R. irregularis* MycFOLD proteins show similarity to chitin-binding CfAvr4 effectors, carbohydrate-binding motif 14 (CBM14) and other carbohydrate-binding domains, whereas their C-termini show similarity to GOLD domains and carry signatures of diversifying selection. Many of the secreted *R. irregularis* MycFOLD proteins are elevated in their expression during colonisation. We furthermore demonstrate that *MycFOLD-17* transcripts accumulate specifically in arbuscules supporting the hypothesis that FOLD proteins act as effectors during plant colonisation by symbiotic fungi.

## Results

### Structural modelling reveals similarities between fungal pathogen effectors and proteins from an AM fungus

Effectors are hallmarks of plant pathogen colonisation success and have also been described from arbuscular mycorrhiza fungi. To address whether the proteome of the AM fungus *R. irregularis* DAOM197198 (Yildirir *et al*., 2021) encodes proteins with structural resemblance to pathogen effectors we carried out a multi-step bioinformatics approach (**Fig. S1**).

A total of 1132 *R. irregularis* proteins were predicted to carry a signal peptide using SignalP 5.0. We excluded all proteins with transmembrane domains predicted via TMHMM 2.0 as well as proteins carrying an endoplasmic reticulum retention signal resulting in 753 candidate extracellular proteins. (**Dataset 1**). Secreted fungal proteins are often further processed by fungal Kexin proteases to give rise to the mature protein version (Outram *et al*., 2021). In our dataset 123 of the 753 proteins have a predicted Kex2 cleavage site following their signal peptide (**Dataset 1**). We then used AlphaFold2 to predict structures of all 753 candidate secreted proteins, after removing predicted signal peptides and Kex2-cleaved pro-domains. From each protein, we used the highest confidence model based on the pLDDT score to establish a structure similarity matrix (**Dataset A, 2-1**) where all protein structures are compared to each other using US-align (Zhang *et al*., 2022). We also included 15 crystal structures and 9 predicted structural models of known effectors from pathogenic fungi, as well as the crystal structure of the DELD family effector Dld1 (PIIN_05872, PDB identifier: 5LOS) from the endophytic fungus *Serendipita indica* (Akum *et al*., 2015; Nostadt *et al*., 2020) and the predicted structural model of RiSLM (Genbank: XM_025315479) from *R. irregularis* (**Table S1**; **Dataset B**; Zeng *et al*., 2020). Next, we established a structural similarity network based on a Template modelling (TM)-score ≥ 0.5 normalized for each pair of protein structures and used the Louvain community detection algorithm (Blondel *et al*., 2008) to extract the communities with similar structures within this network (**Fig. 1**).

**Figure 1.**
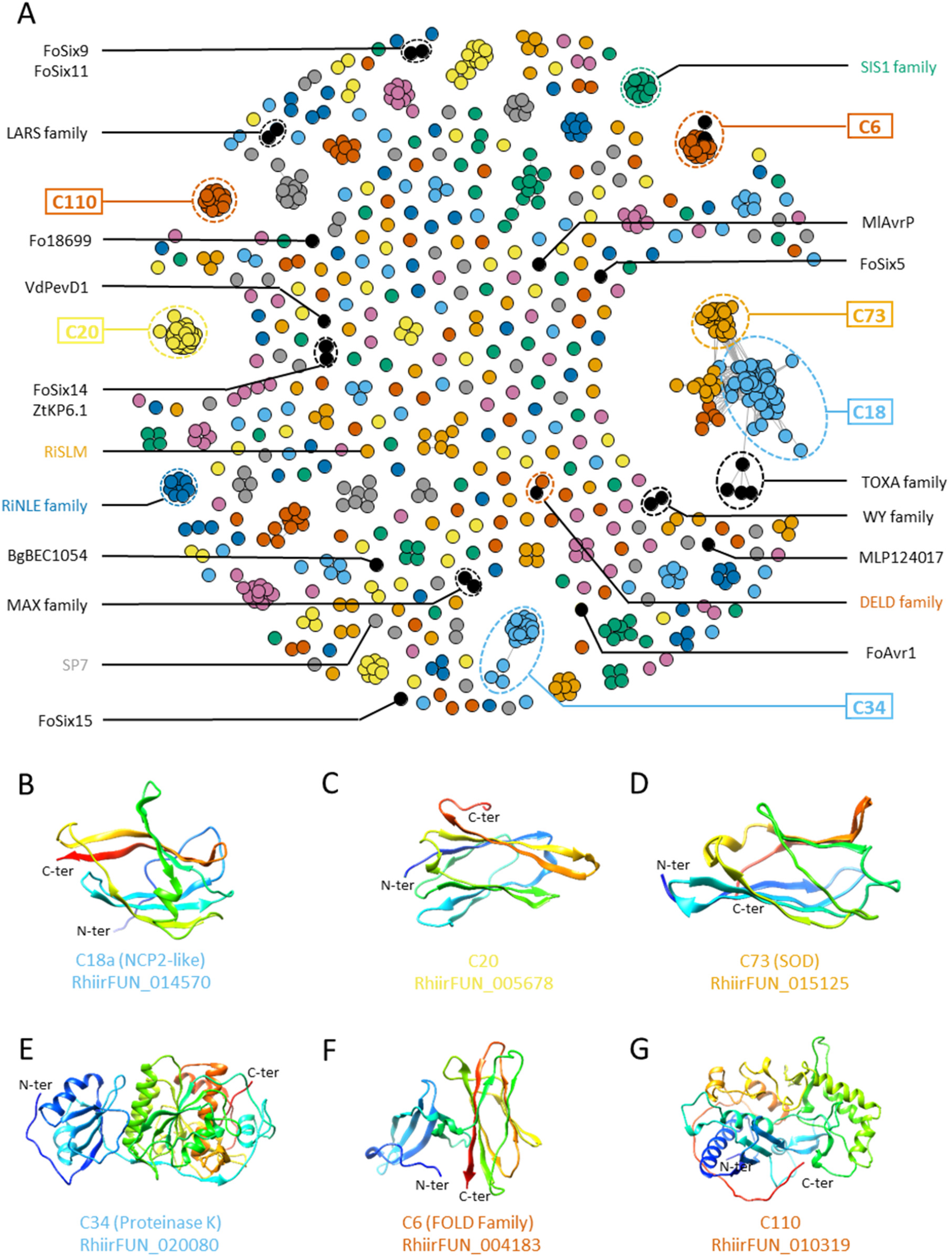
Structural similarities between secreted *R. irregularis* proteins and fungal pathogen effectors. **(A)** Structural similarity network and community analysis of AlphaFold2-modelled secreted *R. irregularis* proteins with previously structurally characterised effectors from oomycete and fungal pathogens (black), an endophyte (red) and known AM fungal effectors (grey). The similarity threshold for clustering was a TM-score of 0.5. **(B-G)** Representative structural models of members of the six largest communities (boxed labels in (A)).

We identified 370 structural communities including 354 only represented by *R. irregularis* proteins (**Dataset 3-1**). The largest community (C18) has 73 members distributed between two sub-communities C18a (possible cholesterol-binding proteins), and C18b (similar to superoxide dismutases) (**Dataset 3-2**). The *R. irregularis* effectors SIS1 and RiNLE1 are part of structural communities C53 (8 members) and C40 (6 members), respectively. The *R*.*irregularis* effectors RiSLM and SP7 are structural singletons (**Fig. 1, Fig. S2, Datasets 1, 3-1**).

Pathogen effectors were only part of one *R. irregularis* community. The pathogen effectors FoSix6 and FoAvr3 were part of the fifth biggest community C6 otherwise represented only by *R*.*irregularis* protein models (**Fig. 1**). In addition, we found a *R. irregularis* protein (RirrFUN_016164) grouping with the *Serendipita indica* DELD effector Dld1 (**Fig. 1, Fig. S2**, Akum *et al*., 2015, Nostadt *et al*., 2020). Given that no other pathogen effectors included in this study shared a community with *R. irregularis* proteins, we centred subsequent analyses on the structural model community C6 harbouring symbiotic *R. irregularis* and *Fol* pathogen proteins.

### *R. irregularis* encodes a large family of FOLD effector candidates

Seventeen *R. irregularis* protein models, FoSix6, and FoAvr3 share a high structural similarity (overall root mean square deviation of atomic positions (RMSD): 1.399; **Fig. 2**) and have a characteristic dual domain configuration. The crystal structure of FoAvr3 was recently elucidated and classified as a FOLD effector (*Fusarium oxysporum* f. sp. *lycopersici* dual domain) together with FoSix6 (Yu *et al*., 2022). We therefore classify these *R. irregularis* protein models as members of the mycorrhiza FOLD (MycFOLD) family.

**Figure 2.**
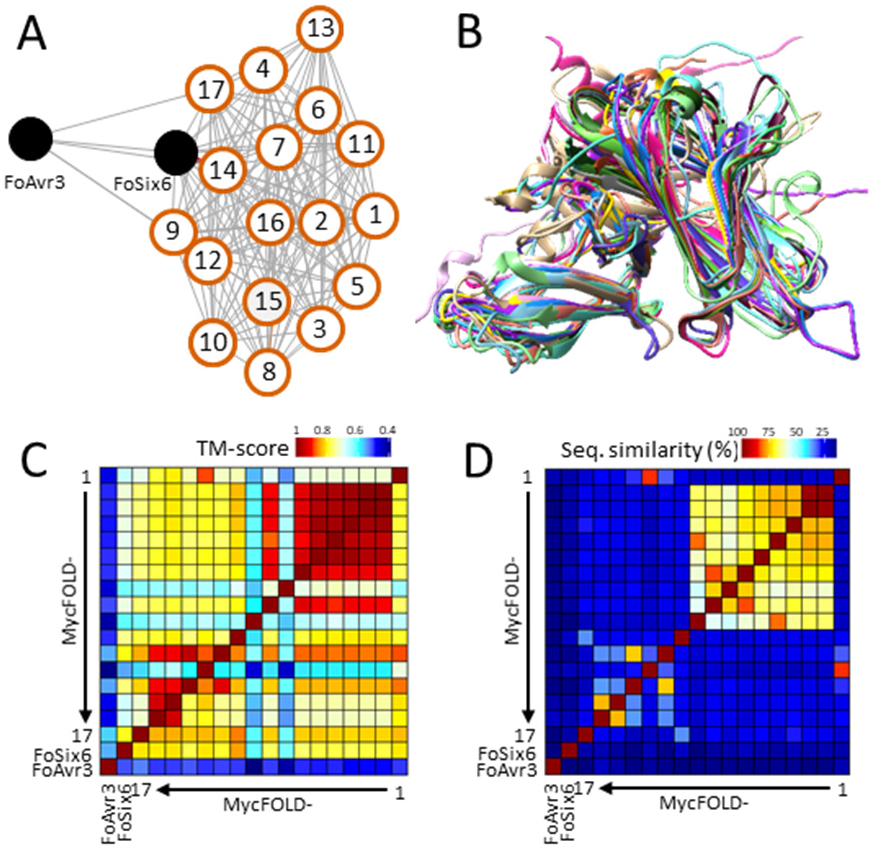
*F. oxysporum* Avr3 and Six6 are structurally but not sequence similar to several *R. irregularis* proteins. Two effectors from *Fol* and 17 *R. irregularis* proteins (numbered) of the community C6 (A) with similar predicted structures (superimposed in B). Models display a high degree of structural similarity reflected in high TM-scores (C) but limited percent sequence similarity (D).

While all MycFOLD members belong to the same structural group (**Fig. 2A-C**), their overall sequence similarity is low (**Fig. 2D**) with exception of 8 highly conserved cysteine residues which form four disulfide bridges (**Fig. S3**). Kex2 cleavage sites reported from FoSix6 and FoAvr3 are predicted in six MycFOLD members (MycFOLD1, −12, −14 to −17) and vary in their position relative to the N-terminus (**Figs. S3 and S4**).

We next derived a hidden Markov model from a sequence alignment of all MycFOLDs as well as FoSix6 and FoAvr3 and surveyed the *R. irregularis* DAOM197198 proteome for additional members of this family. This revealed an additional 9 members which were not part of the secretome as their gene models do not encode signal peptides or both domains of the FOLD structure (**Dataset 4-1**). Together, the *R. irregularis* DAOM197198 genome encodes 26 proteins (17 secreted, nine non-secreted proteins) with a predicted structural similarity to the Six6 FOLD structure.

### MycFOLD proteins combine a predicted carbohydrate binding motif with a diversified Golgi Dynamics (GOLD) domain

FOLD proteins consist of two domains linked together (Yu *et al*., 2022). A protein structural database survey of each domain (**Fig. 3, Database 5**) revealed that the N-terminal domain is similar to carbohydrate-binding module 14 (CBM14) present in dust mite allergen proteins (PDB identifier: 2MFK) and CfAvr4 (PDB identifier: 6BN0, Hurlburt *et al*., 2018).

**Figure 3.**
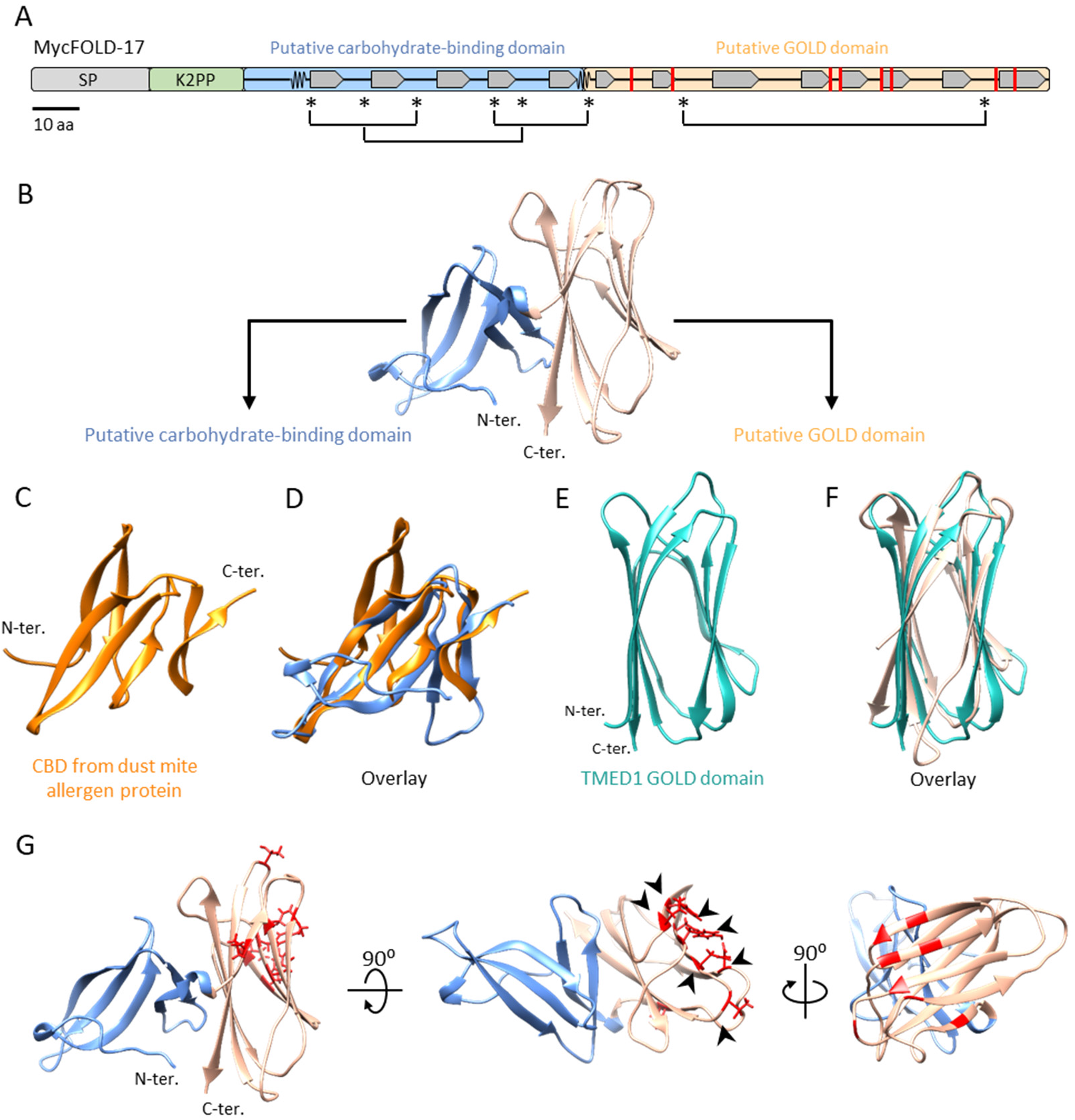
MycFOLD proteins share structural similarities with carbohydrate binding motifs and GOLD domains. (A) MycFOLD proteins have a signal peptide (SP), often a Kex2-processed pro-domain (K2PP) followed by a typical *F. oxysporum* f. sp *lycopersici* dual-domain (FOLD) structure. Conserved and closely positioned cysteine residues likely forming disulphide bridges are indicated with asterisks (A). The N-terminal domain displays structural similarity to the chitin-binding domain from dust mite allergen proteins; RMSD: 2.21; TM-score: 0.501 (C,D), the C-terminal domain structurally matches the GOLD domain of TMED1; RMSD: 2.57; TM-score: 0.684 (E,F). Amino acid residues under highest probability for diversifying selection (arrows) among *R. irregularis* DAOM197198 MycFOLDs are all located on one patch of the second domain (G).

The C-terminal domain of most MycFOLD protein models frequently resembles the Golgi Dynamics (GOLD)-domain found in several eukaryotic Golgi and lipid-traffic proteins such as Transmembrane EMP24 domain containing protein (TMED1, 11/17 MycFOLD models) involved in early endomembrane vesicle trafficking processes (PDB identifier: 7RRM, Mota *et al*., 2022). A disulphide bridge typical of GOLD domains is also predicted for MycFOLDs between two well conserved cysteines (**Fig. S3**). Previously characterised GOLD domains often co-occur with lipid, sterol or fatty acid-binding domains, whereas those in MycFOLDs are coupled with a potential carbohydrate-binding module.

Through a phylogenetic analysis we found patterns of diversifying selection among the MycFOLDs within the *R. irregularis* DAOM197198 genome. Residues with the highest probability for diversifying selection are all located in the C-terminal domain which structurally resembles a GOLD domain (**Fig. 3G, S4**).

### MycFOLD genes are distributed across the genome

Mapping the location of MycFOLD genes showed their distribution across different chromosomes of the *R. irregularis* DAOM197198 genome, where they are present mostly as singletons (**Fig. S5A**). An exception is chromosome 6 which harbours a total of 11 MycFOLD genes, two singletons and a cluster of nine MycFOLD genes. All genes in this cluster (*MycFOLD-2* to *10*) likely evolved from a common ancestor (**Fig. S5B-C**) suggesting that this cluster arose from a recent gene family expansion event.

### MycFOLD transcripts are increased during AM fungal colonisation and accumulate in arbuscules

To address whether MycFOLD genes are transcriptionally upregulated during root colonisation we generated RNA-seq data from *R. irregularis* spores and compared these to data from *Nicotiana benthamiana* colonisation (Dallaire *et al*., 2021) and other publicly available datasets of *M. truncatula* and *B. distachyon* colonisation (Kamel *et al*., 2017; Rich *et al*., 2017; Zeng *et al*., 2018). Our analysis suggests that several MycFOLD transcripts are significantly upregulated during colonisation in all three tested plant species (*MycFOLD-2, −3, −4, −9, −16, −17*) while others are only upregulated during colonisation of two species or even only in *N. benthamiana* (*MycFOLD-8, −10, −11, −12*) suggesting host-specificity of expression (**Fig. 4A; Dataset 6-1**). *MycFOLD-12* showed reduced transcript levels in the *N. benthamiana* dataset. We independently confirmed the higher expression of several MycFOLD genes as well as the absence of a transcript induction of *MycFOLD-12* during *N. benthamiana* root colonisation using qRT-PCR (**Fig. S6**). We did not find any expression support for *MycFOLD-13* or *MycFOLD-15* across colonisation experiments in five different plant species (**Fig. 4A-B; Dataset 6**).

**Figure 4.**
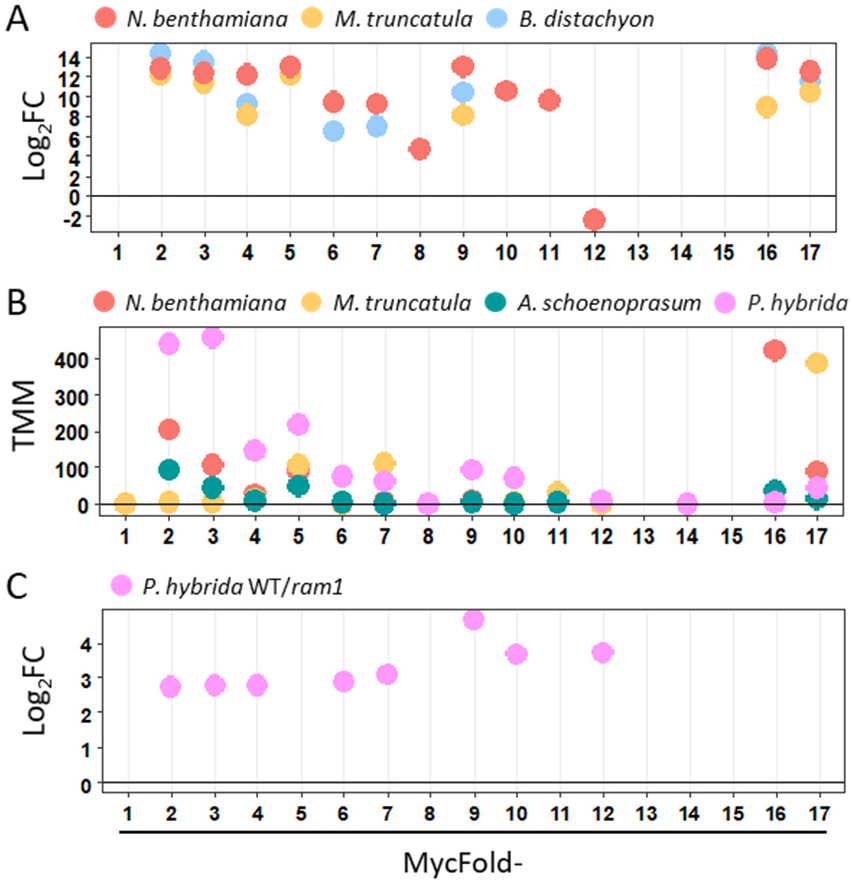
MycFOLD genes show varying patterns of transcriptional upregulation in different host plants. RNA-sequencing analysis of MycFOLD gene expression in datasets from *R. irregularis* comparing (A) colonisation of *N. benthamiana* (Dallaire *et al*., 2021), *Medicago truncatula* and *Brachypodium distachyon* with the corresponding spore-only expression data (Kamel *et al*., 2017); (B) *N. benthamiana, M. truncatula, Allium schoenoprasum* and *Petunia hybrida (*Rich *et al*., *2017*; Zeng *et al*., 2018) trimmed mean-of-M (TMM) levels; and (C) increased MycFOLD transcript levels in wild type *Petunia hybrida* compared to *ram1* mutant (Rich *et al*., 2017).

Comparing the MycFOLD gene expression in *Petunia* wild type and *ram1* plants suggests that eight MycFOLD genes are higher expressed in the wild type. *Petunia ram1* mutants do not form fully branched arbuscules (Rich *et al*., 2015; Rich *et al*., 2017). Expression of some MycFOLD genes may thus be associated with arbuscule development or function (Fig. 4C).

To address the spatial expression of MycFOLD genes in *R. irregularis* colonisation structures in *M. truncatula* roots, we carried out *in situ* hybridisation with *MycFOLD-17* because it is expressed in all analysed plants (**Fig. 4**). We obtained an antisense probe-specific label within arbuscules while the control sense probe did not yield a signal. Counterstaining with wheat germ agglutinin (WGA) coupled to Alexa Fluor 488 showed that almost all arbuscules accumulate *MycFOLD-17* transcript whereas there was no widespread signal at fungal hyphae (**Fig. 5**).

**Figure 5.**
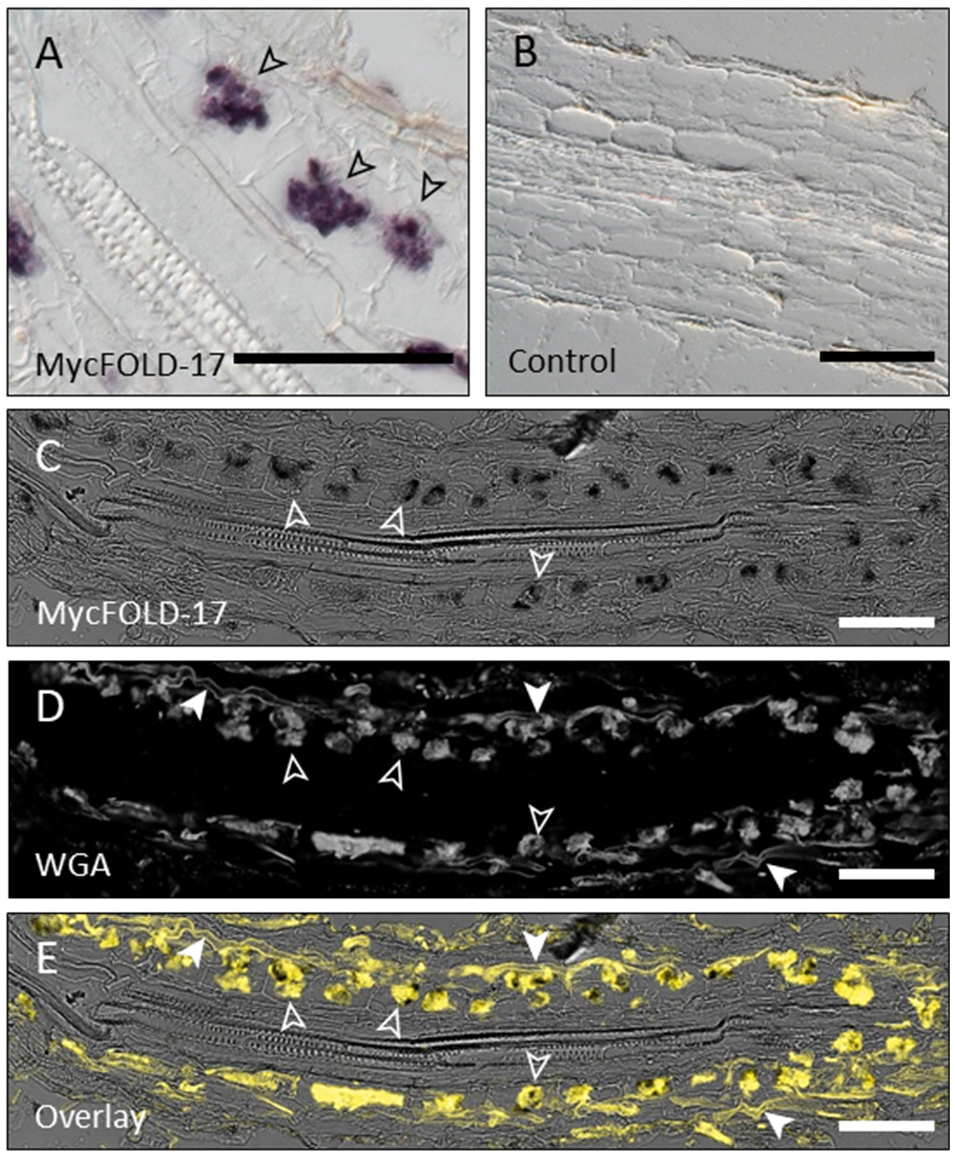
*MycFOLD-17* transcripts accumulate in arbuscules during *Medicago trunctatula* colonisation. *In situ* hybridisation with a *MycFOLD-17* probe (A, C) or control (B). (D) Fungal chitin counterstaining with Alexa Fluor 488-labelled wheat germ agglutinin (WGA). (E) overlay of (C) and (D). Open arrows indicate arbuscules, filled arrows point to intercellular hyphae. Scale bar is 100 μm.

### FOLD proteins are present only in plant-colonising fungi

The structure-based identification of FOLD proteins allowed us to establish a Hidden Markov model (HMM) derived from the 17 secreted *R. irregularis* MycFOLDs and the pathogen FOLD proteins FoAvr3, FoSix6 and thus better encompassing their diversity on the sequence level. We used this MycFOLD-HMM to survey more than 2000 fungal proteomes of the JGI Mycocosm collection (**Dataset 4-2, 4-3**; Grigoriev *et al*., 2014) for presence and abundance of FOLD proteins and predicted the protein structures of the 454 identified matches (**Dataset C**). We then clustered all 454 MycFOLD-HMM matching structure models into communities together with our previously identified *R. irregularis* MycFOLDs, FoSix6 and FoAvr3 (**Fig. S7, Dataset 2-2, 3-3**). Through this we found 388 possible FOLD proteins across 48 fungal species (**Dataset 4-1**). Interestingly, FOLD proteins seem to be only present in fungi which are known to interact with plants. We found extensive variation in the presence and number of matches across the fungal kingdom (**Fig. 6A, Dataset 4-1**). Plant pathogens have a low number of FOLD genes ranging from one to three. By contrast, some Glomeromycotina fungi have much higher FOLD gene numbers, in particular *Diversispora epigaea* and *R. irregularis* strain C3 both with 26 secreted MycFOLD proteins. Notably, some fungi including species of *Funnelliformis* and *Ambispora* did not return any gene matches. In *Paraglomus* and *Diversispora* groups we found sister species with several or no MycFOLD genes.

**Figure 6.**
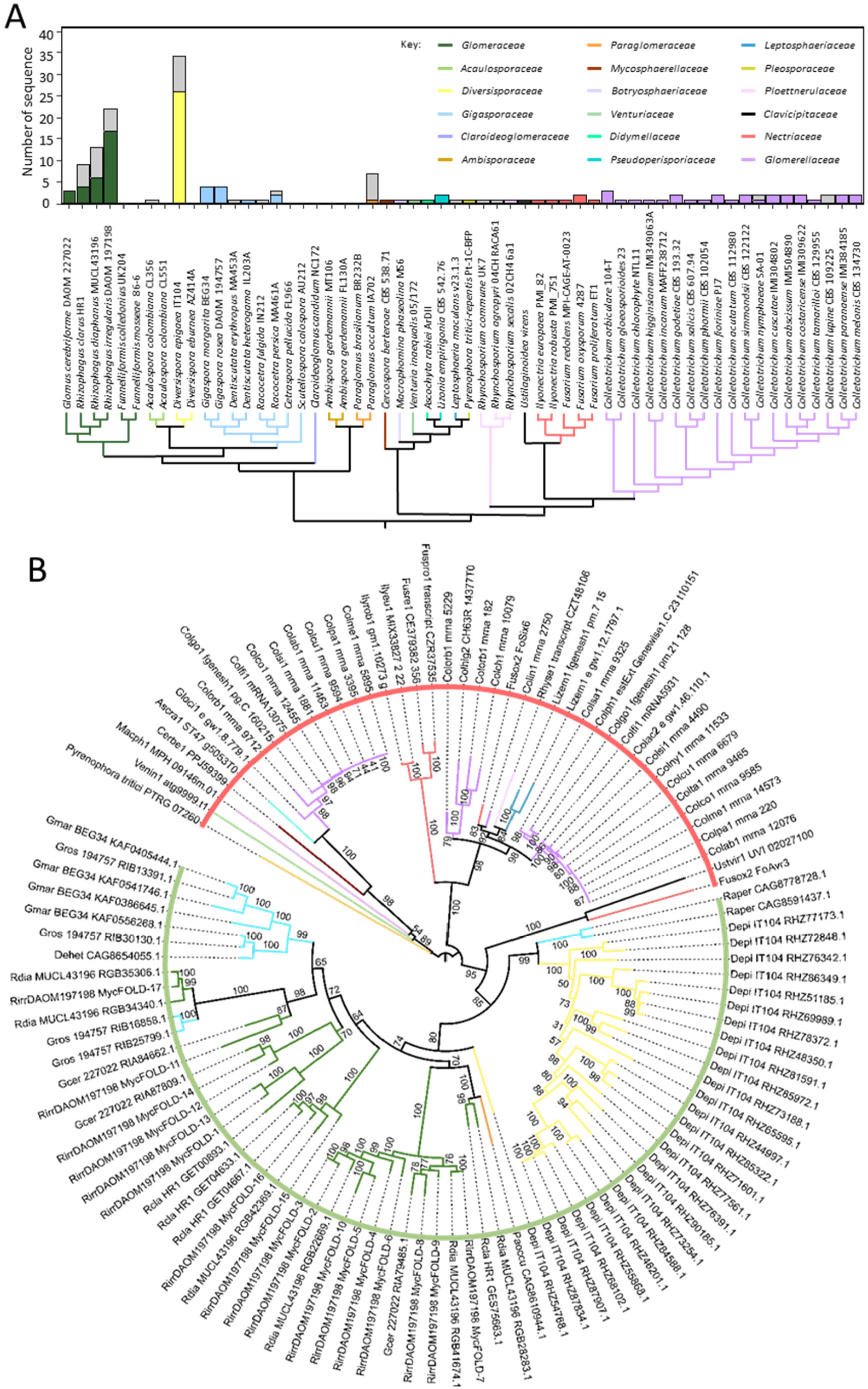
Number and diversity of FOLD genes in fungi. (A) Fungal cladogram with numbers of secreted FOLD genes (colored) and additional non-secreted FOLD genes (gray). One strain per fungal species is displayed. Plant pathogenic fungi genomes encode up to three FOLD proteins, while some Glomeromycotina genomes have up to 30 copies. (B) Phylogenetic analysis of all identified secreted FOLD proteins in proteomes of representative symbiotic (green line) and pathogenic (red line) representative fungal species.

A phylogenetic analysis showed a clear separation between MycFOLDs from Glomeromycotina fungi and FOLD proteins from other plant-interacting fungi (**Fig. 6B**). Interestingly, there are clade specific phylogroups with proliferations and diversification of gene copies, a prominent example is *D. epigaea*.

## Discussion

We used a systematic protein structure modelling approach to classify the secretome of the AM fungus *Rhizophagus irregularis* DAOM197198. We identified secreted proteins with structural similarity to *Fusarium oxysporum* f. sp. *lycopersici* FOLD effectors, which occur in low numbers in fungal pathogen genomes. By contrast, MycFOLD proteins from AM fungi belong to enlarged and diversified gene families when present. We and others (Yu *et al*., 2022) found secreted FOLD proteins across the fungal kingdom. However, they seem to be limited to fungi which interact with plants raising the possibility that they support plant colonisation processes.

Our phylogenetic analysis suggests that MycFOLDs and a subset of pathogen genome encoded FOLDs share a common ancestry but no repeated horizontal gene transfer between pathogenic and symbiotic fungi is evident. It is thus likely that the FOLD configuration was recruited once into Glomeromycotina fungi giving rise to symbiosis specific MycFOLDs. A further dynamic resulting in their loss or proliferation in some AM fungal lineages is evident. By contrast to AM fungal genomes, FOLD gene numbers in plant pathogens remained low, possibly due to their avirulence activities. The FOLD proteins Avr3 (Six1) and Avr1 (Six4) are both recognised by cognate plant membrane receptors (Rep *et al*., 2005; Houterman *et al*., 2008) in shoot tissues. Plant pathogens retain FOLD effectors because they are required for full pathogenicity or to suppress effector-triggered immunity (Houterman *et al*., 2008; Gawehns *et al*., 2014). It will be interesting to test, whether MycFOLDs of symbiotic fungi have the potential to be recognised by the plant immune system in shoots or roots.

FOLD effectors in pathogens and symbiotic fungi consist of two domains. Their individual functions are not well delineated but both might have carbohydrate binding activity. The N-terminal domain of FOLD proteins shows striking similarity to the carbohydrate-binding motif (CBM14) and other effectors which have previously been shown to bind to chitin (CfAvr4, van den Burg *et al*., 2007) or pectin (*Pseudocercospora fuligena* PfAvr4-2, Chen *et al*., 2021). Our analysis revealed structural similarity to GOLD domains including a conserved disulphide bridge. GOLD domains are present in diverse proteins, but a combination with a recognisable CBM14 domain seems limited to FOLD proteins. GOLD domains share significant similarities with sugar-binding proteins, mainly with their CBMs and TMED1 GOLD domains have the potential to oligomerise (Mendes and Costa-Filho, 2022). The C-terminal domains of *R. irregularis* DAOM197198 MycFOLDs display signatures of diversifying selection, and this could point to variable binding specificities for this protein family. Given that secreted FOLD proteins are only present in plant-interacting fungi it is tempting to speculate that they bind to a range of plant carbohydrates possibly at the host-fungal interface.

*MycFOLD* genes are differentially expressed in colonisation of different host plants (**Fig. 4; Dataset 6**). This matches previous observations that a small subset of secreted *R. irregularis* proteins are differentially expressed during colonisation in a host-specific manner. The underlying cues for this host-dependent expression remain unknown (Zeng *et al*., 2018). We found that *MycFOLD-17* transcripts labelled by an RNA probe exclusively accumulate in arbuscules, similar to other AM fungal effectors (e.g., RiNLE1; Wang *et al*., 2021). Whether all MycFOLD genes are expressed at arbuscules remains to be tested. Strikingly, expression levels of MycFOLD genes drop in Petunia *ram1* mutants which are impaired in arbuscule development.

In conclusion, predicting the structures of the *R. irregularis* secretome and comparing this dataset with pathogen effector structures and models has revealed a large family of FOLD proteins previously well characterised as effectors of fungal pathogens. Their exclusive presence in plant-interacting fungi across the fungal kingdom, their inducible expression, lineage specific sequence diversification, and the observed expression at arbuscules support the hypothesis that MycFOLD proteins act as effectors facilitating plant colonisation.

## Materials and Methods

### Biological material and growth condition

Seedlings of *N. benthamiana* were surface sterilised in a solution of 70% ethanol and 0.05% SDS for 3min and washed 3 times in sterile water. Seeds were sown on solid ½ Murashige and Skoog medium (Murashige and Skoog, 1962) and kept at 24C with 16h of photoperiod. After 2 weeks, plants were transferred on silver sand and inoculated with approximately 3200 spores or with water for mock controls. Plants were then kept in a grow chamber (21C the day, 19C the night, 16h photoperiod). Plants were watered with milliQ water every second day for two weeks and fertilized once per week with Long Ashton Solution (Timoneda *et al*., 2021) and milliQ water on the other days. Colonized roots were collected 4 weeks after inoculation for subsequent analysis.

*R. irregularis* strain DAOM197198 was co-cultivated from monoxenic carrot (*Daucus carota*) root organ culture on solid M Medium (Becard and Fortin, 1988) in the dark at 25C.

#### Spore harvesting and germination

Spores were harvested from monoxenic carrot root organ culture of *R. irregularis* strain DAOM197198 by dissolving the phytagel in citrate buffer (8.3mM sodium citrate dihydrate, 1.7mM citric acid, pH 6) for 20min under stir. The solution was first filtered through muslin cloth and then through a 45um pores metal sieve. Spores were washed twise in sterile water and their concentration was determined using a Malassez counting chamber.

For spore germination, approximatively 60,000 spores were incubated at 30C with 2% CO2 for 7 days in liquid M-medium. Germinated spores were collected using a 40-um cell strainer (Sigma-Aldrich).

#### Quantitative PCR

Total RNAs from germinated spores and roots of *N. benthamiana* colonized by *R. irregularis* were extracted using the Qiagen RNeasy Mini Kit (Qiagen) and treated with DNase I (Qiagen). Three biological replicates were performed per sample. Height hundred nanograms of the total RNAs per sample were reverse transcribed using iScript cDNA Synthesis Kit (Biorad). Real time quantitative PCR were performed using LightCycler 480 SYBR Green I Master Mix (Roche) and the transcript levels of MycFOLD-2, 9, 12, 16 and 17 genes were normalized to *R. irregularis* elongation factor 1α (RiEF-1α) and β-tubulin (RiβTub) using 34 cycles as threshold. Primers used to amplify MycFOLD transcripts were listed in table S2.

#### Secretome prediction

The most recent version of the *R. irregularis* annotation (Yildirir *et al*., 2022) was used for the secretome prediction. All 26,820 annotated proteins were analysed by SignalP 5.0 (Almagro-Armenteros *et al*., 2019) for the signal peptide prediction. Secreted candidates carrying a transmembrane domain predicted by TMHMM v2.0 (Krogh *et al*., 2001), as well as an endoplasmic reticulum retention motif [K/H]DEL (Pelham, 1990) at the C-terminus, were removed. Mature proteins of less than 15 amino acids length were removed. The functional annotation of the 753 secreted candidates was performed using the annotate function of the Funannotate pipeline (version 1.8.13; Palmer *et al*., 2020).

#### Structure model prediction

The protein structure predictions were performed using the 753 secreted proteins and MycFOLD-HMM hits after removal of the predicted signal peptides. For each protein, the presence of a kexin protease cleavage site was predicted as described previously (Outram *et al*., 2021) and when multiple cleavage sites were predicted, the KEX2 prodomain with the higher probability was removed. *ab initio* prediction was carried out using ParaFold (Zhong *et al*., 2021) which allows large-scale structure predictions with AlphaFold2 (Jumper *et al*., 2021). The multiple sequence alignment (msa) step was running on the full databases. Five models were generated for each protein. The structural model predictions of all proteins are available (Teulet & Schornack, 2022; 10.5281/zenodo.7443325). The best model (ranked_0) was determined by the predicted LDDT (pLDDT) scores.

Because of the gene coding for the recently described RiSLM effector (Zeng *et al*., 2020) was not annotated in the *R. irregularis* genomic annotation used in this study, its structural prediction was determined apart from the other secreted proteins by Alphafold2 using the ColabFold online tool (Mirdita *et al*., 2022) and the best model determined by the pLDDT score was selected.

#### Structural similarity search

For each secreted protein model, we used Foldseek (van Kempen *et al*., 2022) to identify the best template from the PDB100 database. The top results with a TM-score over 0.5 were selected.

#### Structural network

Structural similarities for all Alphafold models, including the 25 additional protein structures from known pathogenic effector families (see **Table S1**), were determined by US-align (Zhang *et al*., 2022) based on the normalized TM-score. Briefly, each structure model was aligned against the others and a matrix of the pair-wise TM-score was built. Structural similarity was considered as significant only for a normalized TM-score ≥ 0.5 in the secretome network and a TM-score ≥ 0.6 in the MycFOLD-HMM hit network, for both structures. Based on this threshold, the TM-score matrix was then converted to an adjacency matrix in which 1 or 0 replace the TM-scores when both protein structures are considered as similar or not, respectively. The adjacency matrix was then turned into a network using the Igraph R package (https://igraph.org/r/) and protein clustering was performed by the Louvain community detection method (Blondel *et al*., 2008) implemented in Igraph.

#### Phylogenetic analysis

All protein alignments were performed using T-Coffee (Notredame *et al*., 2000). The phylogeny was inferred from the resulting alignment using the Maximum Likelihood method as implemented in IQ-TREE (Nguyen *et al*., 2014) and branch supports were estimated after 10,000 iterations of ultrafast bootstrap. The resulting consensus trees were then used for subsequent analysis. Signatures of diversifying selection of MycFOLDs were determined by PAMLX (Xu and Zhang, 2013) using the CodeML module. First, a codon alignment of the 17 MycFOLD coding sequences was performed by PAL2NAL (Suyama *et al*., 2006) guided by a protein sequence alignment. The codon alignment, and its resulting phylogenetic tree built by IQ-TREE as described above, were then used to run CodeML Model 3 (discrete) with default parameter and fix ω = 0.

#### Sequence analysis

FOLD sequence homology searches were carried out on two protein sequences databases, the entire non-redundant fungal proteomic database from JGI mycocosm (more than 2000 proteomes, released in July 2021; https://mycocosm.jgi.doe.gov/mycocosm/home; **Dataset 4-2**; Grigoriev *et al*., 2014) and a database built from all available proteomes of 36 strains of Glomeromycotina (**Dataset 4-3**). A protein sequence alignment of all the proteins belonging to the FOLD family cluster C6 was used as input for the Hidden Markov Model (HMM) research of FOLD homologs. An HMM profile was first built by *hmmbuild* (HMMER package v3.3.2, Eddy, 2011) and then used as input in the *hmmsearch* program to identify new FOLD homologs in the two fungal databases mentioned above with a score > 20 as threshold. The structural models and clustering of the 454 output sequences were then subsequently analysed as described above for the *R. irregularis* secretome.

#### RNA in-situ hybridization of wax-embedded root sections

*Medicago truncatula* roots colonised with *R. irregularis* were fixed in Formalin-Aceto-Alcohol (FAA), then dehydrated and embedded in paraffin using a Leica ASP300 tissue processor. Paraffin moulds were prepared with a Leica EG1160 paraffin embedding station. A Leica X microtome was used to obtain 10 μm-thick sections. Slides with longitudinal root sections were selected for *in situ* hybridization. A 666 bp fragment of MycFOLD-17 cDNA was amplified using the gene-specific primers MF17_ISH_F (ATGAGAATATTTTCGGCTCAA) and MF17_ISH_R (TCAGTCAGTTATCCAAGCAGC) and ligated into the pGEM-T Easy vector (Promega) and used as a template for *in vitro* transcription using the Digoxygenin (DIG) RNA Labeling Kit (Roche) and SP6 RNA polymerase. T7 RNA polymerase was used to generate the sense probe used as negative control. Wax was removed from sections in HistoChoice (Sigma) and root samples were rehydrated prior to treatment with proteinase K. After dehydration, the samples were incubated with antisense or control sense RNA probes overnight at 55°C. Hybridised probes were detected with anti-DIG antibodies and revealed with alkaline phosphatase-conjugated antibodies in presence of nitro blue tetrazolium (NBT) and 5-bromo-4-chloro-3-indolyl phosphate (BCIP, Roche).

#### WGA staining

*R. irregularis* cell walls were stained with Wheat Germ Agglutinin (WGA) conjugated with Alexa Fluor 488 (Thermo) using a protocol adapted from Bonfante-Fasolo *et al*., (1990). Briefly, root sections were incubated in a staining solution (25 μg/ml WGA in phosphate-buffered saline (PBS)) for 2 hours in the dark, washed in PBS, mounted in 10% glycerol and imaged with a Leica SP8 confocal microscope (Leica, Wetzlar, Germany) equipped with a 60× NA 1.2 water immersion objective. An excitation wavelength of 488 nm was used for imaging. The *in-situ* hybridization signal was detected through transmitted light.

#### RNAseq analysis

Fastq files (**Table S3**) were downloaded from the NCBI Biosample database and quality checked by FastQC (https://www.bioinformatics.babraham.ac.uk/projects/fastqc/). When applicable, overrepresented sequences were removed by Trimmomatic (Bolger *et al*., 2014). The filtered reads were then aligned on the *R. irregularis* genome (Manley *et al*., 2022) by STAR (v2.5.3a modified) using default parameters. Raw counts were obtained by FeatureCount from the Rsubread package (Liao *et al*., 2019) with default parameters and only uniquely mapped reads were considered further. Differential expression analyses between colonised root and germinated spore samples were carried out by Bioconductor-DESeq package (Love *et al*., 2014) on R. The absolute log2 fold change (Log2FC) was used to visualise the differential MycFOLD transcript levels. Trimmed mean-of-M (TMM) normalisations were obtained using the Bioconductor-EdgeR package (Robinson *et al*., 2010) from raw count tables generated by FeatureCount. The visualised data represent the TMM average of the three biological replicates.

## Acknowledgements

The authors would like to thank Alexandra Dallaire Gokalp Yildirir and Nicolas Corradi for providing revised gene model predictions for *R. irregularis*, Francois Nedelec for providing a Python script to streamline AlphaFold batch prediction. We also are grateful to Philip Carella and Alex Guyon for critical comments on the manuscript.

## Author contributions

AT, CQ, EE, WY and SS conceived and designed experiments. AT, CQ, EE, AW, WY and SS performed experiments and data analyses. AT, EE, AW, and SS wrote the manuscript.

## Supplementary Figures

**Figure S1. A multi-step bioinformatics approach to identify AM fungal proteins with structural similarity to fungal plant pathogen effectors**. SP, signal peptide; TM, transmembrane domain; ER, endoplasmic reticulum; K2PP, Kex2-processed pro-domain; TM-score, Template-modelling score.

**Figure S2. Predicted *R. irregularis* communities with structural similarity**. (A) Community cluster sizes. (B) Sub-communities within community C18 based on a TM-score ≥ 0.6. (C-H) Representative structural models from communities C18a, C18b, C53, C40, C369 and C14, the latter displaying similarity to DELD effectors Dld1 from *Serendipita indica*. Colouring follows a rainbow gradient from N to C-terminus. TM-score, Template modelling score.

**Figure S3. MycFOLD proteins have shared protein features with FoSix6 and FoAvr3**. (A) Protein sequence alignment with signal peptides (yellow highlight), a predicted KEX2 protease processing site (green highlight) and conserved cysteines (asterisks). (B-D) location of the disulfide bonds (yellow) in the structure models of MycFOLD-17, FoSix6, and FoAvr3. The domains of the FOLD fold are coloured differently.

**Figure S4. Patterns of diversifying selection in *R. irregularis* MycFOLDs**. The barplot along the MycFOLD protein alignment represents the Naive Empirical Bayes probabilities of each site for a dN/dS ratio (omega) > 3.961, estimated by codeml model M3 (discrete). Blue and red bars represent posterior probabilities *p* < 0.95 and *p* < 0.99, respectively. The N-terminal and C-terminal protein domains are underlined in blue and beige, respectively. The conserved cysteine residues are indicated by an asterisk. The signal peptides are highlighted in yellow and the predicted KEX2 protease processing site in green.

**Figure S5. MycFOLD genes are distributed across the *R. irregularis* genome**. (A) Subgenomic locations of MycFOLD genes. (B) Phylogenetic analysis of the clustered MycFOLD genes suggests they share a common ancestor. (C) Detailed view of the gene cluster on chromosome 6, MycFOLD genes are red, other genes white.

**Figure S6. Validation of MycFOLD gene induction in colonised *N. benthamiana* roots (Myc) and spores**. Gene expression of selected genes was quantified using quantitative reverse transcriptase PCR. All gene expression values were normalized using *R. irregularis* elongation factor 1 alpha (*RiEF-1α*) and beta-tubulin (*RiβTub*). The mean +/-standard deviation (black) of three biological replicates (gray) are plotted. Grey dots represent actual values. One-Way ANOVA was used in statistical analysis.

**Figure S7. Structurally similar communities of MycFOLD HMM hits**. (A) Community clusters and their size based on the normalised TM-score ≥ 0.6. The structural communities with FOLD similarity are indicated. (B) Structural similarity network and community analysis of each AlphaFold2-modelled MycFOLD-HMM hits. (C-K) Representative structural models from FOLD/MycFOLD communities C1, C4, C2, C26, C13, C20, C21, C23 and C55. The N-terminal and C-terminal protein domains are underlined in blue and beige, respectively.

## Supplementary Tables

**Table S1. Additional structural models of effector proteins used in the study**. This includes the protein and effector family names, the PDB structure identifier when available, as well as the community number to which the protein belongs in the secretome network. For the proteins without PDB identifier, an Alphafold2 prediction model is used for the Louvain community detection.

**Table S2. RT-qPCR primers used in the study**.

**Table S3. Sequence Read Archive (SRA) accession number used for RNAseq analysis in the study**. This includes the Bioproject as well as the SRA numbers of raw read samples used for the *de novo* mapping.

## Supplementary Datasets

**Dataset 1 - Genomic and structural features of the secreted proteins used in this study**. This includes genomic location, coding strand, gene name, functional annotation via Funannotate, SignalP 5.0 and Kex2-processed pro-domain (K2PP) cleavage site predictions, and the structural features of the best model for each proteins including the predicted LDDT (pLDDT) score, the Louvain community cluster to which the protein belongs in the secretome structural network and the results of the Foldseek analysis with the best template identified based on the TM-score. Furthermore, DNA, CDS and translated protein sequences for each gene, including the mature proteins after removal of signal peptide or K2PP when applicable are listed.

**Dataset 2 - US-align TM-scores of pairwise structural comparisons of highest-confidence AlphaFold2 models of all proteins used in the study**. (2-1) All TM-scores resulting from the pairwise structural comparisons done by US-align of each structural model compared to the others. (2-2) All TM-scores resulting from the pairwise structural comparisons done by US-align of each MycFOLD-HMM hit compared to the others.

**Dataset 3 - Structural community composition analyses**. Communities identified by the Louvain community detection method are sorted by community size starting with the largest community. When possible, annotations are provided. (3-1) *R. irregularis* secretome network communities; (3-2) Secretome sub-community C18; (3-3) Communities from MycFOLD-HMM hits across the fungal kingdom including those which have no predicted overall similarity to FOLD proteins.

**Dataset 4 - Genomic and structural features of the MycFOLD-HMM hits identified in fungal proteomic databases**. (4-1) Structural features of each MycFOLD-HMM hit from the JGI and Glomeromycotina fungal proteome databases including SignalP 5.0 and Kex2-processed prodomain (K2PP) cleavage site predictions, structural features of the best model for each protein and their predicted LDDT score, and the Louvain community cluster to which the protein belongs in the MycFOLD structural network. Furthermore, protein sequences for each MycFOLD-HMM hits, including the mature proteins after removal of signal peptide or K2PP when applicable are listed (4-2) Biological information and NCBI accession numbers of fungal genomes from the JGI database (4-3) GenBank accession numbers and additional information for all Glomeromycotina proteomes used in this study

**Dataset 5 - DALI-lite database matches of FOLD N- and C-terminal domains**. Five best matches identified for the N- and C- terminal of the 17 MycFOLDs, FoSix6 and FoAVR3 effectors present in the Protein Data Bank.

**Dataset 6 - Raw transcript counts of reanalysed RNAseq experiments**. Gene IDs from gene annotation provided by Manley et al., 2022 (a) and Yildirir et al., 2021 (b). *nd*, not detected. (6-1) RNAseq mapping statistics and differential gene expressions of the 17 *MycFOLD* genes during AM fungal colonisation of various host plant species by *Rhizophagus irrgularis* compared to germinated spores only. (6-2) Trimmed mean-of-M (TMM) values of the 17 *R. irregularis MycFOLD* genes during colonisation of various host plant species.

