## Supplementary Figures and Tables for "A pathogen effector FOLD diversified in symbiotic fungi"

Article title: A pathogen effector FOLD diversified in symbiotic fungi

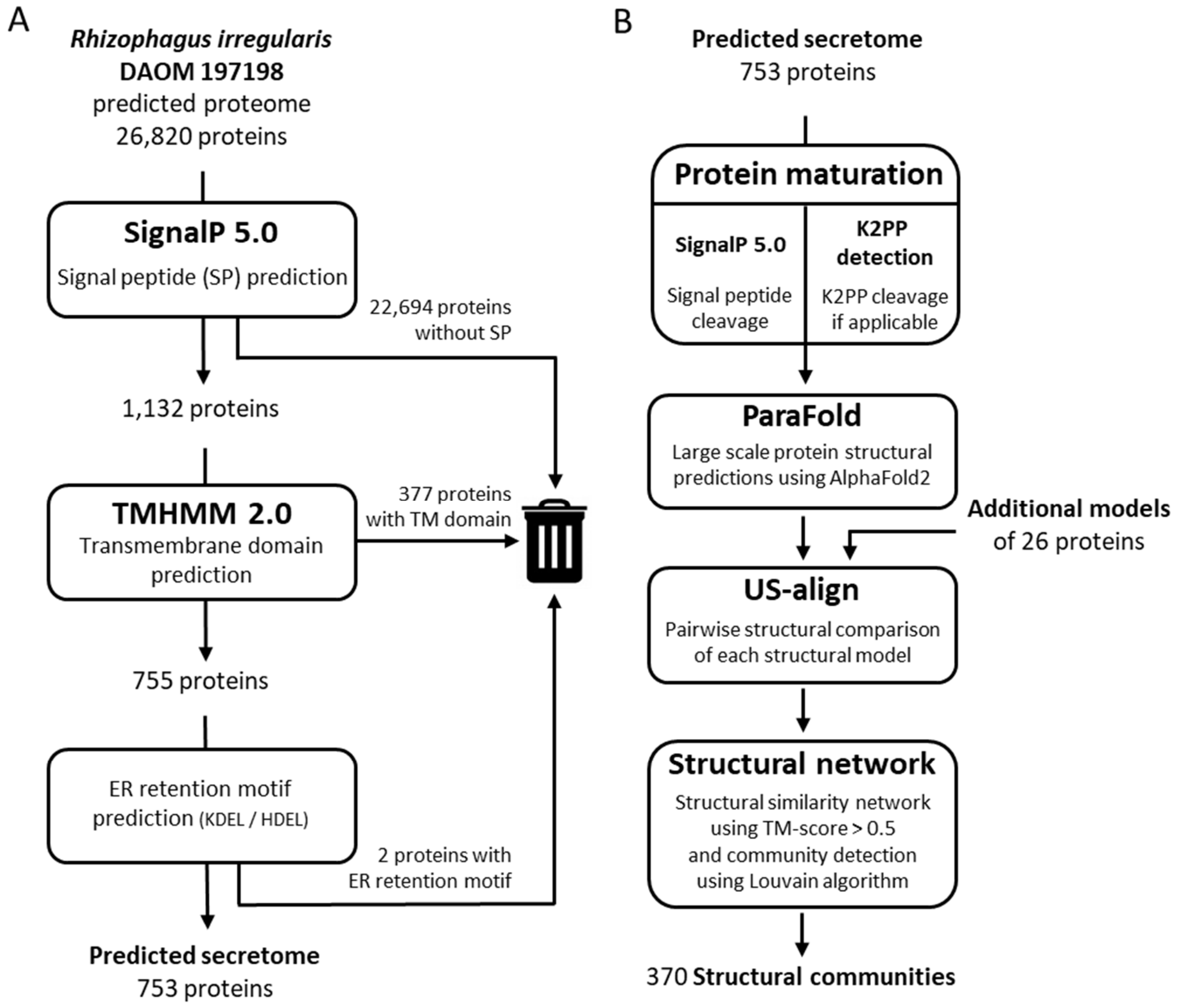


**Figure S1. A multi-step bioinformatics approach to identify AM fungal proteins with structural similarity to fungal plant pathogen effectors.** SP, signal peptide; TM, transmembrane domain; ER, endoplasmic reticulum; K2PP, Kex2-processed pro-domain; TM-score, Template-modelling score.

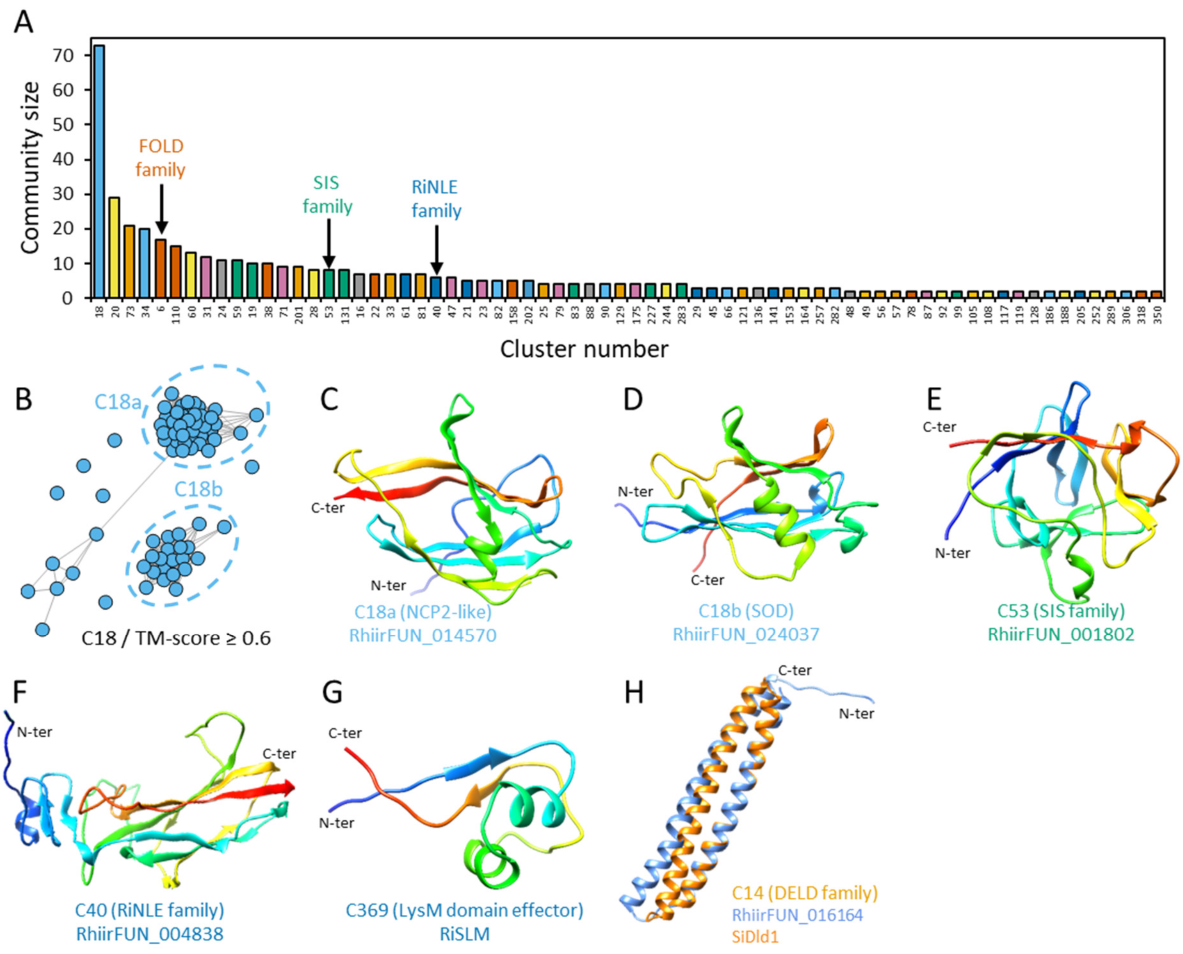


**Figure S2. Predicted *R. irregularis* communities with structural similarity.** (A) Community cluster sizes. (B) Sub-communities within community C18 based on a TM-score ≥ 0.6. (C-H) Representative structural models from communities C18a, C18b, C53, C40, C369, and C14, the latter displaying similarity to DELD effectors Dld1 from *Serendipita indica*. Colouring follows a rainbow gradient from N to C-terminus. TM-score, Template modelling score.

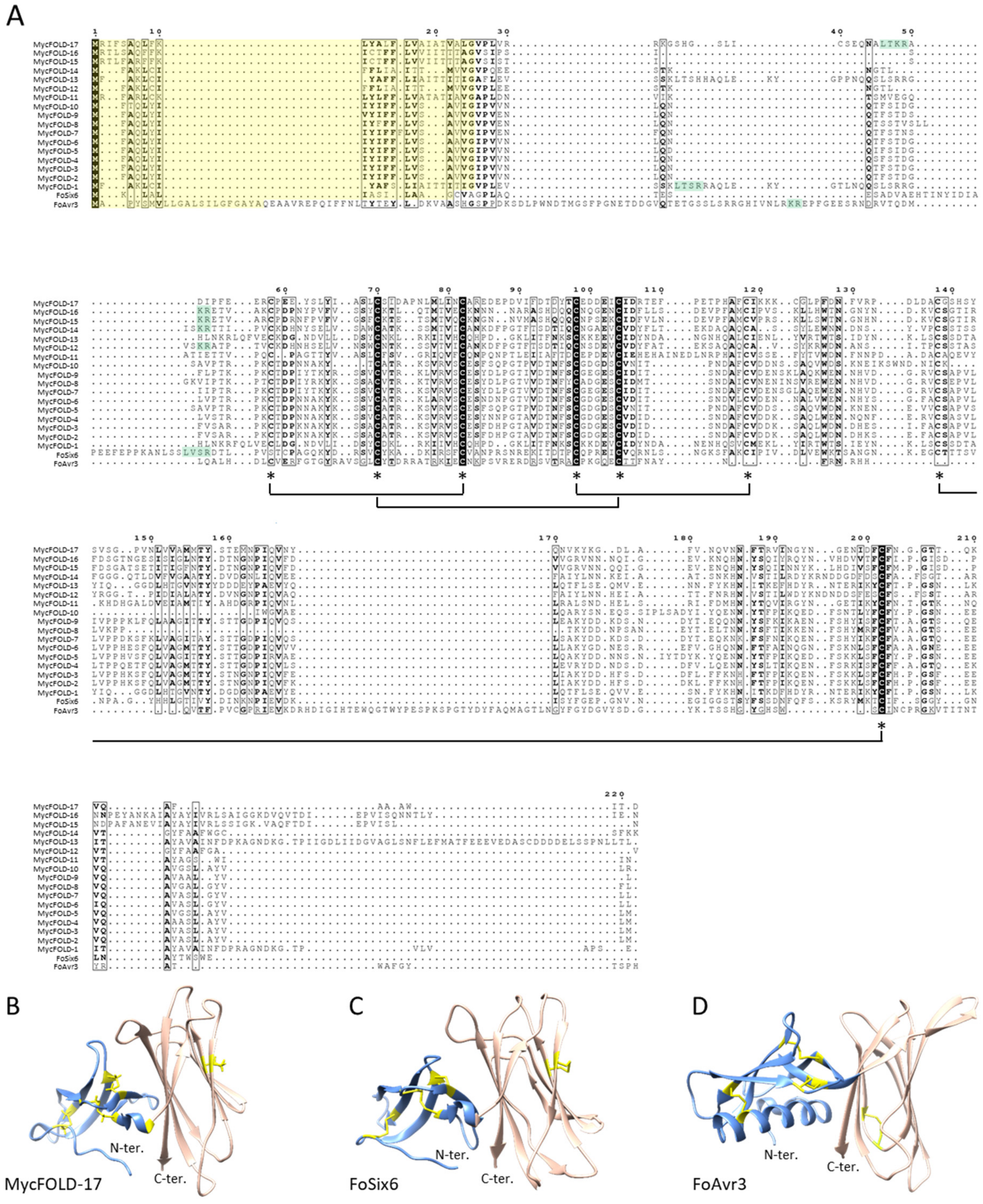


**Figure S3. MycFOLD proteins have shared protein features with FoSix6 and FoAvr3.** (A) Protein sequence alignment with signal peptides (yellow highlight), a predicted KEX2 protease processing site (green highlight) and conserved cysteines (asterisks). (B-D) location of the disulfide bonds (yellow) in the structure models of MycFOLD-17, FoSix6, and FoAvr3. The domains of the FOLD fold are coloured differently.

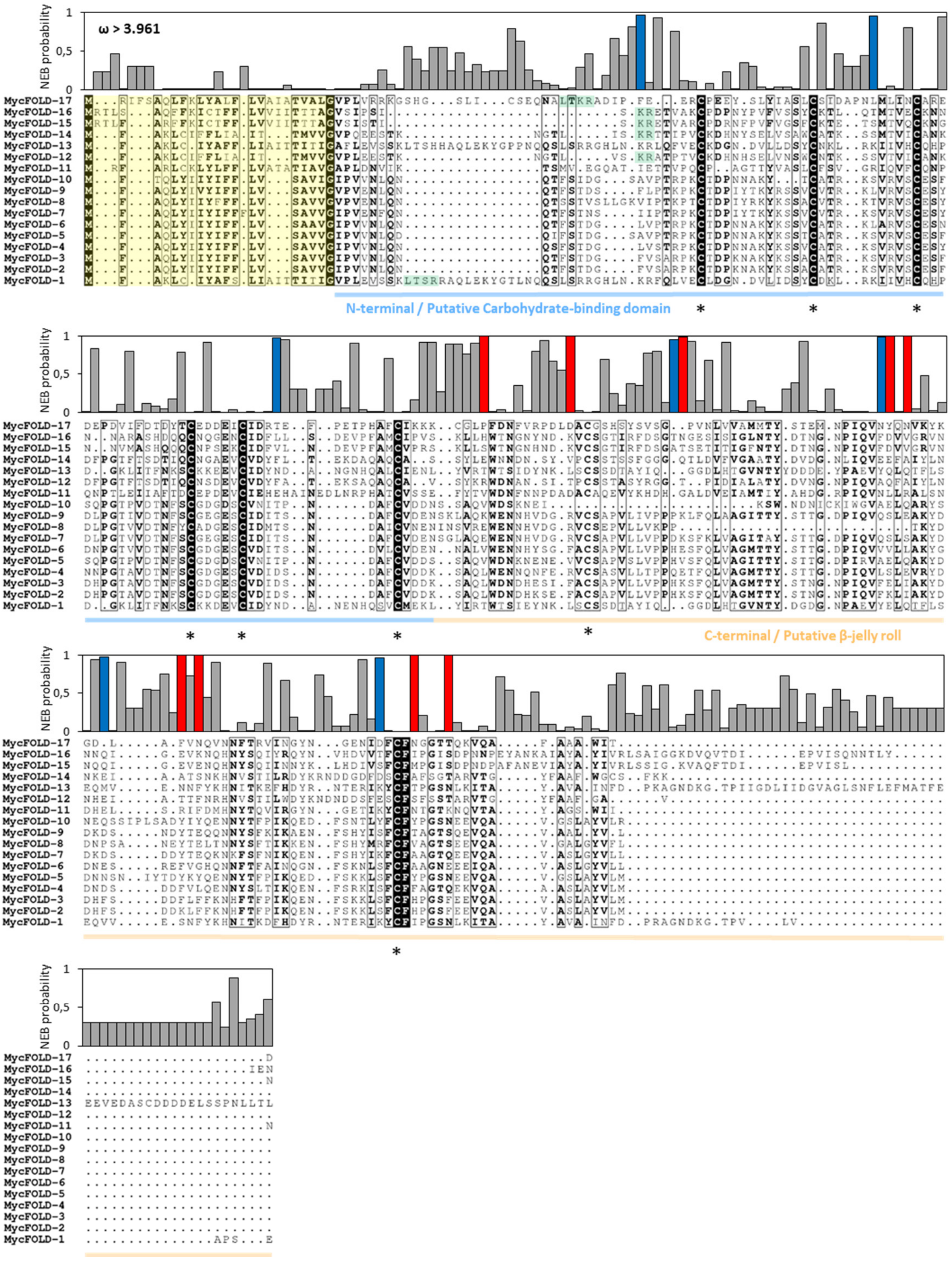


**Figure S4. Patterns of diversifying selection in *R. irregularis* MycFOLDs**. The barplot along the MycFOLD protein alignment represents the Naive Empirical Bayes probabilities of each site for a dN/dS ratio (omega) > 3.961, estimated by codeml model M3 (discrete). Blue and red bars represent posterior probabilities $p < 0.95$ and $p < 0.99$, respectively. The N-terminal and C-terminal protein domains are underlined in blue and beige, respectively. The conserved cysteine residues are indicated by an asterisk. The signal peptides are highlighted in yellow and the predicted KEX2 protease processing site in green.

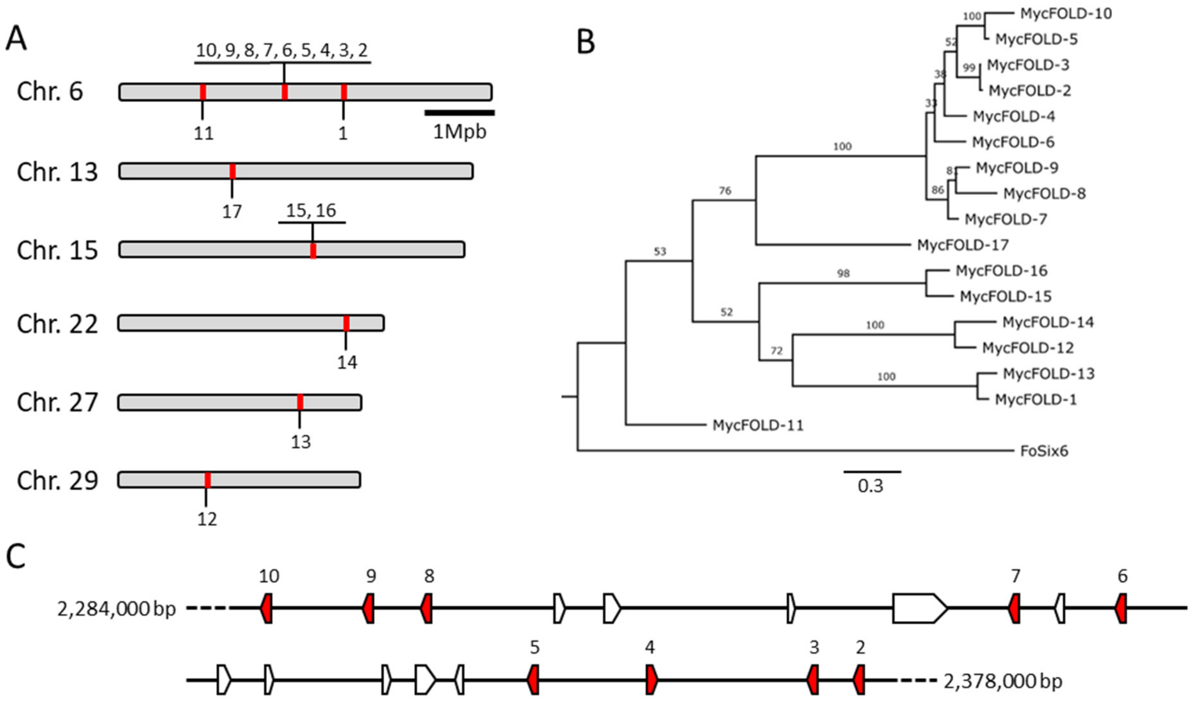


**Figure S5. MycFOLD genes are distributed across the *R. irregularis* genome.** (A) Subgenomic locations of MycFOLD genes. (B) Phylogenetic analysis of the clustered MycFOLD genes suggests they share a common ancestor. (C) Detailed view of the gene cluster on chromosome 6, MycFOLD genes are red, other genes white.

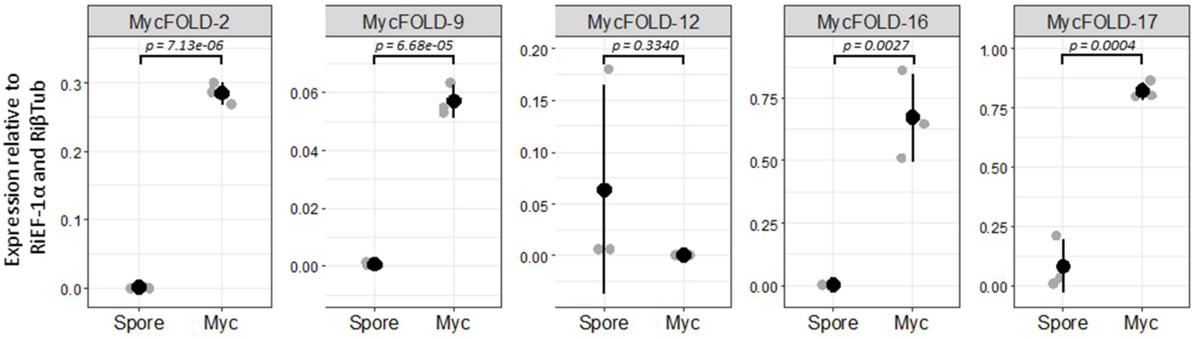


**Figure S6. Validation of *MycFOLD* gene induction in colonised *N. benthamiana* roots (Myc) and spores.** Gene expression of selected genes was quantified using quantitative reverse transcriptase PCR. All gene expression values were normalized using *R. irregularis* elongation factor 1 alpha (*RiEF-1α*) and beta-tubulin (*RiβTub*). The mean +/- standard deviation (black) of three biological replicates (gray) are plotted. Grey dots represent actual values. One-Way ANOVA was used in statistical analysis.

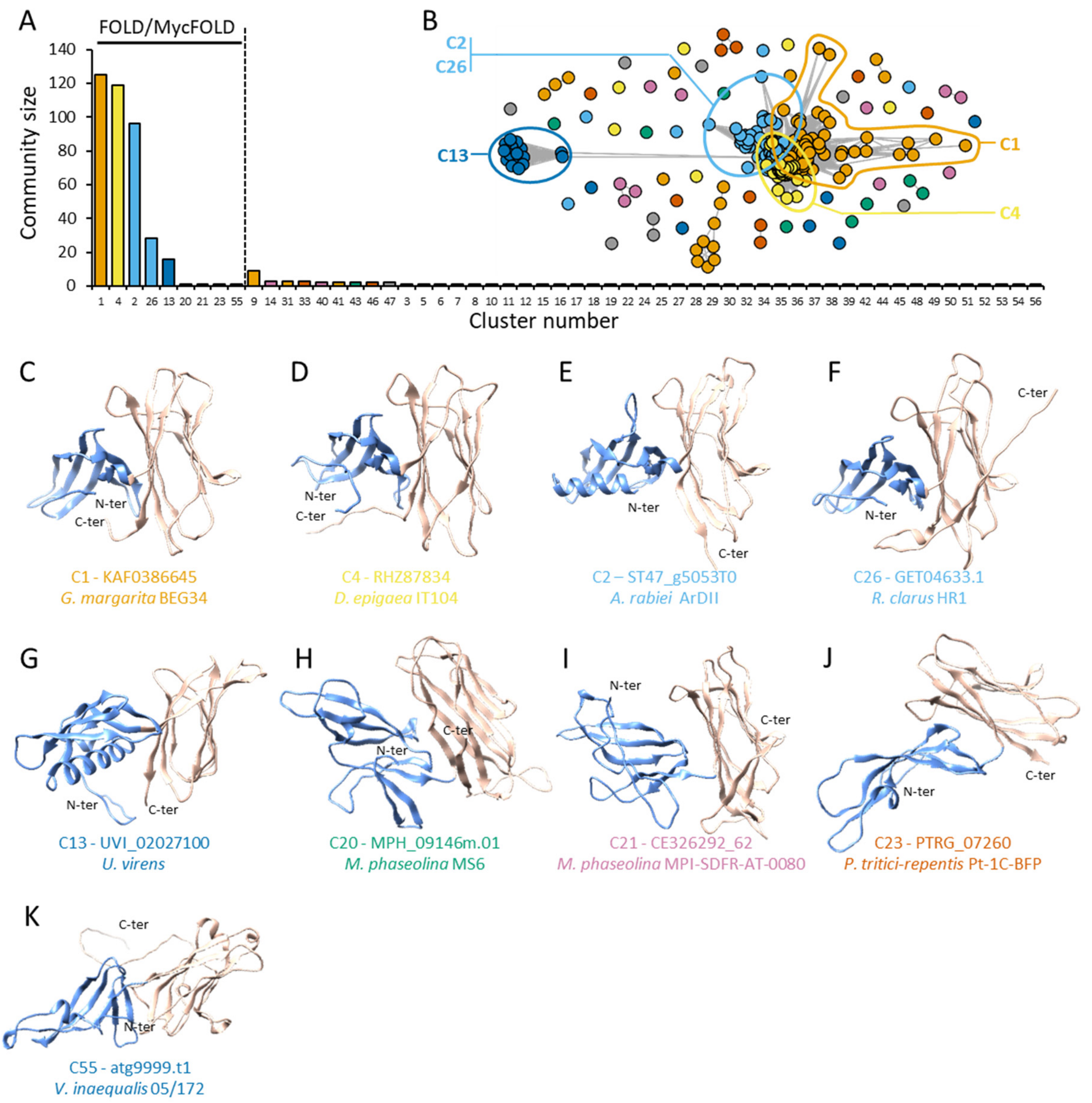


**Figure S7. Structurally similar communities of MycFOLD HMM hits.** (A) Community clusters and their size based on the normalised TM-score ≥ 0.6. The structural communities with FOLD similarity are indicated. (B) Structural similarity network and community analysis of each AlphaFold2-modelled MycFOLD-HMM hits. (C-K) Representative structural models from FOLD/MycFOLD communities C1, C4, C2, C26, C13, C20, C21, C23 and C55. The N-terminal and C-terminal protein domains are underlined in blue and beige, respectively.

| Effector family | Protein name | Organism | PDB identifier | Community number | Source |
|---|---|---|---|---|---|
| ToxA | PtrToxA | *Pyrenophora tritici-repentis* | 1ZLD | C5 | Sarma *et al.*, 2005 |
| | FoAvr2 | *Fusarium oxysporum* f. sp. lycopersici | 5OD4 | C5 | Di et *al.*, 2017 |
| | MlAvrL567-A | *Melampsora lini* | 2OPC | C5 | Guncar *et al.*, 2007 |
| | MlAvrL567-D | *Melampsora lini* | 2QVT | C5 | Wang *et al.*, 2007 |
| WY RXLR | PcAvr3a11 | *Phytophthora capsici* | 3ZR8 | C15 | Boutemy *et al.*, 2011 |
| | PiPexRD2 | *Phytophthora infestans* | 3ZRG | C15 | Boutemy *et al.*, 2011 |
| MAX | MoAvr-Pia | *Magnaporthe oryzae* | 6Q76 | C13 | Varden *et al.*, 2019 |
| | MoAvr1-CO39 | *Magnaporthe oryzae* | 5ZNG | C13 | Guo *et al.*, 2018 |
| RALPH | BgBEC1054 | *Blumeria graminis* | 6FMB | C1 | Pennington *et al.*, 2019 |
| LARS | LmAvrLm5-9 | *Leptosphaeria maculans* | 7AD5 | C2 | Lazar *et al.*, 2022 |
| | FfEcp11-1 | *Fulvia fulva* (*Cladosporium fulvum*) | 6ZUS | C2 | Lazar *et al.*, 2022 |
| FOLD | FoAvr1 | *Fusarium oxysporum* f. sp. lycopersici | Alphafold2 prediction | C4 | This study |
| | FoSix6 | *Fusarium oxysporum* f. sp. lycopersici | Alphafold2 prediction | C6 | This study |
| | FoAvr3 | *Fusarium oxysporum* f. sp. lycopersici | Alphafold2 prediction | C6 | This study |
| Fusarium family 3 | FoSix9 | *Fusarium oxysporum* f. sp. lycopersici | Alphafold2 prediction | C7 | This study |
| | FoSix11 | *Fusarium oxysporum* f. sp. lycopersici | Alphafold2 prediction | C7 | This study |
| Fusarium family 4 | FoSix14 | *Fusarium oxysporum* f. sp. lycopersici | Alphafold2 prediction | C8 | This study |
| | FoSix5 | *Fusarium oxysporum* f. sp. lycopersici | Alphafold2 prediction | C10 | This study |
| Fusarium family 5 | FoSix15 | *Fusarium oxysporum* f. sp. lycopersici | Alphafold2 prediction | C9 | This study |
| | FOXGR_18699 | *Fusarium oxysporum* f. sp. lycopersici | Alphafold2 prediction | C3 | This study |
| LysM | ZtKP6-1 | *Zymoseptoria tritici* | 6QPK | C8 | Padilla *et al.*, 2020 |
| NTF2-like | MLP124017 | *Melampsora larici-populina* | 6SGO | C11 | de Guillen *et al.*, 2019 |
| DELD | SiDld1 | *Serendipita* (*Piriformospora*) *indica* | 5LOS | C14 | Nostadt *et al.*, 2020 |
| Zn-binding | MlAvrP | *Melampsora lini* | 5VJJ | C12 | Zhang *et al.*, 2018 |
| C2-like | VdPevD1 | *Verticillium dahliae* | 5XMZ | C370 | Zhou *et al.*, 2017 |
| LysM | RiSLM | *Rhizophagus irregularis* | Alphafold2 prediction | C369 | This study |

**Table S1. Additional structural models of effector proteins used in the study**. This includes the protein and effector family names, the PDB structure identifier when available, as well as the community number to which the protein belongs in the secretome network. For the proteins without PDB identifier, an Alphafold2 prediction model is used for the Louvain community detection.

| Gene id | Gene name | Primer | Sequence 5' -> 3' |
|---|---|---|---|
| RhiirFUN_023549 | *MycFOLD-2* | forward | CAAATGATGCATTCTGTGTTGACG |
| | | reverse | GTGCTGAACATGCGAATATACTTTC |
| RhiirFUN_023532 | *MycFOLD-9* | forward | AGTGCCACCACCAAAGTTAT |
| | | reverse | TCTAGCGATTGTACTTGTATTGGA |
| RhiirFUN_017017 | *MycFOLD-12* | forward | AGCGACGAAGTTTGTGTTGATT |
| | | reverse | GCATTGTCCCATCTCTTGTACG |
| RhiirFUN_006262 | *MycFOLD-16* | forward | CAGACCCCAACTATCCTGTGT |
| | | reverse | TCGTGACTAGCCCTTGCATT |
| RhiirFUN_004183 | *MycFOLD-17* | forward | CAAGAGAGGATGAGCCAGACG |
| | | reverse | GCATGTGGGGTCTCTGGAAAT |
| RhiirFUN_003909 | *Riβtub* | forward | AACAATTGGGCCAAAGGTCACT |
| | | reverse | CGCTTCTTTGCGAACAACATCT |
| RhiirFUN_009362 | *RiEF-1α* | forward | TGTTGCTTTCGTCCCAATATC |
| | | reverse | GGTTTATCGGTAGGTCGAG |

**Table S2. RT-qPCR primers used in the study.**

| Source | BioProject | Plant partner | Fungal partner | Sample | SRA accession |
|---|---|---|---|---|---|
| Dallaire *et al.*, 2021 | PRJNA722386 | *Nicotiana benthamina* | *Rhizophagus irregularis* DAOM 197198 | Nb + Ri rep1 | SRR14294955 |
| | | | | Nb + Ri rep2 | SRR14294954 |
| | | | | Nb + Ri rep3 | SRR14294953 |
| | | - | *Rhizophagus irregularis* DAOM 197198 | Germinated spore rep1 | SRR14294961 |
| | | | | Germinated spore rep2 | SRR14294960 |
| | | | | Germinated spore rep3 | SRR14294959 |
| Kamel *et al.*, 2017 | PRJNA281452 | *Medicago truncatula* | *Rhizophagus irregularis* DAOM 197198 | Mt + Ri rep1 | SRR1979303 |
| | | | | Mt + Ri rep2 | SRR1979304 |
| | | | | Mt + Ri rep3 | SRR1979305 |
| | | *Brachypodium distachyon* | *Rhizophagus irregularis* DAOM 197198 | Bd + Ri rep1 | SRR1979306 |
| | | | | Bd + Ri rep2 | SRR1979307 |
| | | | | Bd + Ri rep3 | SRR1979308 |
| | PRJNA209330 | - | *Rhizophagus irregularis* DAOM 197198 | Germinated spore rep1 | SRR3285893 |
| | | | | Germinated spore rep2 | SRR3285894 |
| | | | | Germinated spore rep3 | SRR3285895 |
| Zeng *et al.*, 2018 | PRJNA389248 | *Medicago truncatula* | *Rhizophagus irregularis* DAOM 197198 | Mt + Ri rep1 | SRR5644313 |
| | | | | Mt + Ri rep1 | SRR5644314 |
| | | | | Mt + Ri rep2 | SRR5644315 |
| | | | | Mt + Ri rep2 | SRR5644316 |
| | | | | Mt + Ri rep3 | SRR5644317 |
| | | | | Mt + Ri rep3 | SRR5644318 |
| | | *Allium schoenoprasum* | *Rhizophagus irregularis* DAOM 197198 | As + Ri rep1 | SRR5644319 |
| | | | | As + Ri rep1 | SRR5644320 |
| | | | | As + Ri rep2 | SRR5644321 |
| | | | | As + Ri rep2 | SRR5644322 |
| | | | | As + Ri rep3 | SRR5644323 |
| | | | | As + Ri rep3 | SRR5644324 |
| | | *Nicotiana benthamiana* | *Rhizophagus irregularis* DAOM 197198 | Nb + Ri rep1 | SRR5644325 |
| | | | | Nb + Ri rep1 | SRR5644326 |
| | | | | Nb + Ri rep2 | SRR5644327 |
| | | | | Nb + Ri rep2 | SRR5644328 |
| | | | | Nb + Ri rep3 | SRR5644329 |
| | | | | Nb + Ri rep3 | SRR5644330 |
| Rich *et al.*, 2017 | PRJNA380056 | *Petunia hybrida* | *Rhizophagus irregularis* DAOM 197198 | Ph + Ri rep1 | SRR5366066 |
| | | | | Nb + Ri rep2 | SRR5366067 |
| | | | | Ph + Ri rep2 | SRR5366068 |
| | | *Phram1-2* mutant | *Rhizophagus irregularis* DAOM 197198 | *ram1* + Ri rep1 | SRR5366072 |
| | | | | *ram1* + Ri rep2 | SRR5366073 |
| | | | | *ram1* + Ri rep3 | SRR5366074 |

**Table S3. Sequence Read Archive (SRA) accession number used for RNAseq analysis in the study**. This includes the Bioproject as well as the SRA numbers of raw read samples used for the *de novo* mapping.
